# *Helicobacter* chronic stage inflammation undergoes fluctuations that are altered in *tlpA* mutants

**DOI:** 10.1101/2022.08.01.502425

**Authors:** Kevin S. Johnson, Christina Yang, J. Elliot Carter, Atesh K. Worthington, Elektra K. Robinson, Raymond Lopez-Magaña, Frida Salgado, Isabelle Arnold, Karen M. Ottemann

## Abstract

*Helicobacter pylori* colonizes half of the world’s population and is responsible for a significant disease burden by causing gastritis, peptic ulcers, and gastric cancer. The development of host inflammation drives these diseases, but there are still open questions in the field about how *H. pylori* controls this process. We characterized *H. pylori* inflammation using an eight month mouse infection time course and comparison of wild type and a previously identified mutant lacking the TlpA chemoreceptor that causes elevated inflammation. Our work shows that *H. pylori* chronic stage corpus inflammation undergoes surprising fluctuations, with changes in Th17 and eosinophil numbers. The *H. pylori tlpA* mutant changed the inflammation temporal characteristics, resulting in different inflammation from wild type at some time points. tlpA mutants have equivalent total and gland colonization at late stage infections. During early infection, in contrast, they show elevated gland and total colonization compared to WT. Our results suggest the chronic inflammation setting is dynamic, and may be influenced by colonization properties of early infection.

**Importance:** *Helicobacter pylori* established chronic infection in half of the world’s population. This infection initiates during childhood, and leads to later life gastric diseases including gastritis, peptic ulcers, and gastric cancer. These diseases are driven by host inflammation, with more severe inflammation leading to more severe disease. It is not fully understood how *H. pylori* controls inflammation. In this work, we used an *H. pylori* mouse infection model and characterized inflammation and colonization of wild-type *H. pylori* and a mutant that was known to cause elevated inflammation. We found that inflammation in the chronic stage undergoes surprising fluctuations, with related changes in Th17 numbers. *H. pylori* tlpA mutants caused offset inflammation dynamics, suggesting that H. pylori infection underlies this dynamism. Although there were not colonization dynamics at the time of inflammation fluctuations, there were differences between WT and *tlpA* mutant colonization at early time points. Our results suggest the chronic inflammation setting is more dynamic than previously thought and may be influenced by colonization properties of early infection.

## Introduction

Inflammation underlies a widespread set of disease states, resulting from diverse triggers that can be abiotic or microbial. In healthy inflammation, the inflammatory response is triggered and then resolves. Some triggers, however, result in chronic inflammation that does not fully resolve, resulting in lifelong disease. One such trigger is the bacterium *Helicobacter pylori*. This bacterium colonizes individuals during early childhood and maintains a lifelong colonization of the stomach (1, 2). Colonization in the very early childhood period leads to immunological benefits, characterized by the development of T-regulatory cells that in turn decrease the risk of asthma and other pathologies (1, 3, 4). Continued colonization into later life, however, can lead to disease, including gastritis, peptic ulcers, and gastric cancer, ultimately contributing to a significant disease burden throughout the world (3, 5–7). These diseases arise from *H. pylori*-driven stomach inflammation. To be able to control these diseases, it is critical to understand the mechanisms by which this bacterium manipulates host inflammation.

*H. pylori* forms intimate interactions with the mammalian host, colonizing two distinct regions of the stomach called the corpus and antrum. *H. pylori* is found in several locations, including the mucus layer lining the gastric epithelium, attached to gastric epithelial cells, and within gastric glands that invaginate both regions (8–11). By all accounts, the stomach does not appear to be a hospitable niche for colonization. Its high acidity is toxic to *H. pylori* (12) and contents of the stomach are emptied within hours of eating a solid digestible meal (13). In addition, the mucus layer (14, 15) and gastric epithelial cells are renewed in a matter of days, a process which is exacerbated by *H. pylori* colonization (16, 17). Despite these challenges, *H. pylori* chronically colonizes the stomach using a variety of factors. These include virulence factors such VacA and γ-glutamyl-transpeptidase (GTT), which both lead to tolerogenic T-cells responses through effects on myeloid cell populations in the stomach (2, 18); the Cag PAI, which alters host cells signaling, increasing host inflammation and disease severity; adherence factors to facilitate host cell attachment; and urease to buffer the local environment of the bacteria; as well as several other factors (2, 9).

*H. pylori* locates to specific regions in the stomach, in a manner that is dictated by its chemotaxis system (19). Chemotaxis regulates bacterial motility in response to harmful and beneficial signals. These signals are sensed by chemoreceptors and are transduced through a two-component signal transduction system (9, 20). *H. pylori* expresses four chemoreceptors, TlpA, TlpB, TlpC, and TlpD. Much work has been done to determine the sensing profile of these chemoreceptors and their roles *in vivo*. Multiple attractants are sensed that are beneficial for *H. pylori* growth, including arginine, fumarate, and cysteine by TlpA; urea by TlpB; and lactate by TlpC (21–25). Repellents, in contrast, are harmful to the bacteria and include acid sensed by multiple chemoreceptors, autoinducer-2 sensed by TlpB, and reactive oxygen species sensed by TlpD (26–30). The ability to sense signals chemotactically is critical for *in vivo* gland colonization, as chemotaxis and chemoreceptor mutants exhibit significant gastric gland colonization defects *in vivo* (8, 28, 31, 32). Ultimately, chemotaxis maximizes the fitness of the bacteria *in vivo*, as essential metabolites can be sought, and harmful host conditions can be avoided.

*H. pylori* chemoreceptors play unique roles during colonization, likely due to their distinct sensing profiles. Some in *H. pylori* promote colonization levels, while others affect inflammation. This work focusses on TlpA, which is required for inflammation control. Specifically, Δ*tlpA H. pylori* induced significantly more inflammation, measured by histology at 6 months post infection (33). At this time point, the Δ*tlpA* mutant does not differ from wild type in colonization levels, although at earlier time points it does have a mild colonization defect in both single strain and competition infections (34) (35). Intriguingly, the high inflammation phenotype associated with loss of TlpA was observed only at six months post infection but not three (33). These results suggest loss of *tlpA* creates an *H. pylori* strain that has subtly altered host interactions. Given that host inflammation control by *H. pylori* dictates disease severity, this study aims to understand how TlpA modulates host inflammation. (36–38).

Inflammation induced by *H. pylori* is initiated by bacterial interactions with innate immune cells, which in turn drive effector CD4^+^ T-cell differentiation through the production of cytokines (18). Infection recruits myeloid cells to the gastric lamina propria, where they interact with *H. pylori*. These cells include neutrophils, eosinophils, multiple types of dendritic cells (DCs) including CD103^+^ CD11b^+^ DCs, CD103^+^ CD11b^-^ DCs, CD103^-^ CD11b^+^ DCs; and multiple types of monocytes and macrophages including MHCII^-^ monocytes, MHCII^+^ monocytes, MHCII^+^ macrophages (18, 39–41). These innate immune cell populations interact with and/or phagocytose *H. pylori*, leading to the recruitment of CD4+ effector T cells. At the later stages of infection, the *H. pylori* induced inflammation consists of large populations of CD4^+^ effector T-cells, including Th1, Th17, and Tregs (42). Th1 and Th17 cells promote pro-inflammatory responses by producing IFNγ and IL-17a, respectively (36, 43–46). Conversely, Tregs suppress inflammation through the secretion of IL-10 (Arnold et al., 2011; Eaton et al., 2001; Gray et al., 2013; Rolig et al., 2011). Overall, the inflammatory response induced by *H. pylori* consists of a mixture of pro- and anti-inflammatory T cells, as well as multiple types of innate immune cells.

The ability of *H. pylori* to colonize gastric glands is important for several reasons. For one, gland colonization seems to underlie colonization resistance, the property of an initial infection preventing a second one. Specifically, non-chemotactic *H. pylori* do not colonize the glands and develop colonization resistance like WT does (32, 47, 48). Second, *H. pylori* in the glands can interact with gastric Lgr5^+^ stem cells, increasing the proliferation rate of these mammalian cells, leading to hyperplasia (17). Lastly, gland populations may be associated with inflammation. Specifically, non-chemotactic mutants do not colonize glands and induce significantly less inflammation (33, 46). This inflammatory response is associated with a decreased pro-inflammatory Th17 cell response compared to WT *H. pylori* (46). This data suggest that gland colonization may promote a pro-inflammatory Th17 response. Other data show that, in addition, the host T-cell response limits *H. pylori* gland infection, further supporting the connection (47, 49). Currently, it is unknown what role TlpA plays in gland colonization or how any phenotypes relate to host inflammation.

In this work, we aimed to better understand the characteristics of *H. pylori* inflammation by assessing gland and total colonization, as well as inflammation development over an 8-month infection course. We compared wild type *H. pylori* to the *tlpA* mutant, to gain insight into inflammation control. We report the surprising finding that H. pylori chronic stage inflammation is not constant, and instead shows regular fluctuations particularly in the corpus region. The *tlpA* mutant show offset and enhanced inflammation compared to WT. These inflammation variations were associated with changes in Th17 cell density. While there were some co-occurring colonization variations, the most substantial colonization differences between WT and the tlpA mutant occurred in early infection, with *tlpA* required to maintain WT level gland loads during early infection. This work suggests that bacterial properties, influenced by TlpA, affects host inflammation in the corpus by attenuating Th17 cell abundance, potentially through the regulation of early gland colonization.

## Results

### *tlpA* mutants result in high fluctuating inflammation that is corpus specific and offset from WT inflammation

Previous work showed that Δ*tlpA H. pylori* mutants induce high levels of inflammation at 6-months post-infection, but normal levels at 3-months (33). We thus examined the inflammation kinetics of WT and *tlpA* mutants. We carried out two independent, 8-month mouse colonization studies, orally infecting with WT or Δ*tlpA* SS1 GFP^+^ *H. pylori*. Inflammation was assessed by histology of hematoxylin and eosin-stained tissue sections spanning the corpus to antrum. Inflammation grades were assigned by a pathologist (Dr. J. Elliot Carter) assessing the density and distribution of lymphocytic infiltration into the lamina propria of the corpus and antrum separately in blinded samples (33, 35). Inflammation grades were determined for both time courses (Sup. Fig. 1) and presented as averages (Fig. 1).

**Figure 1.**
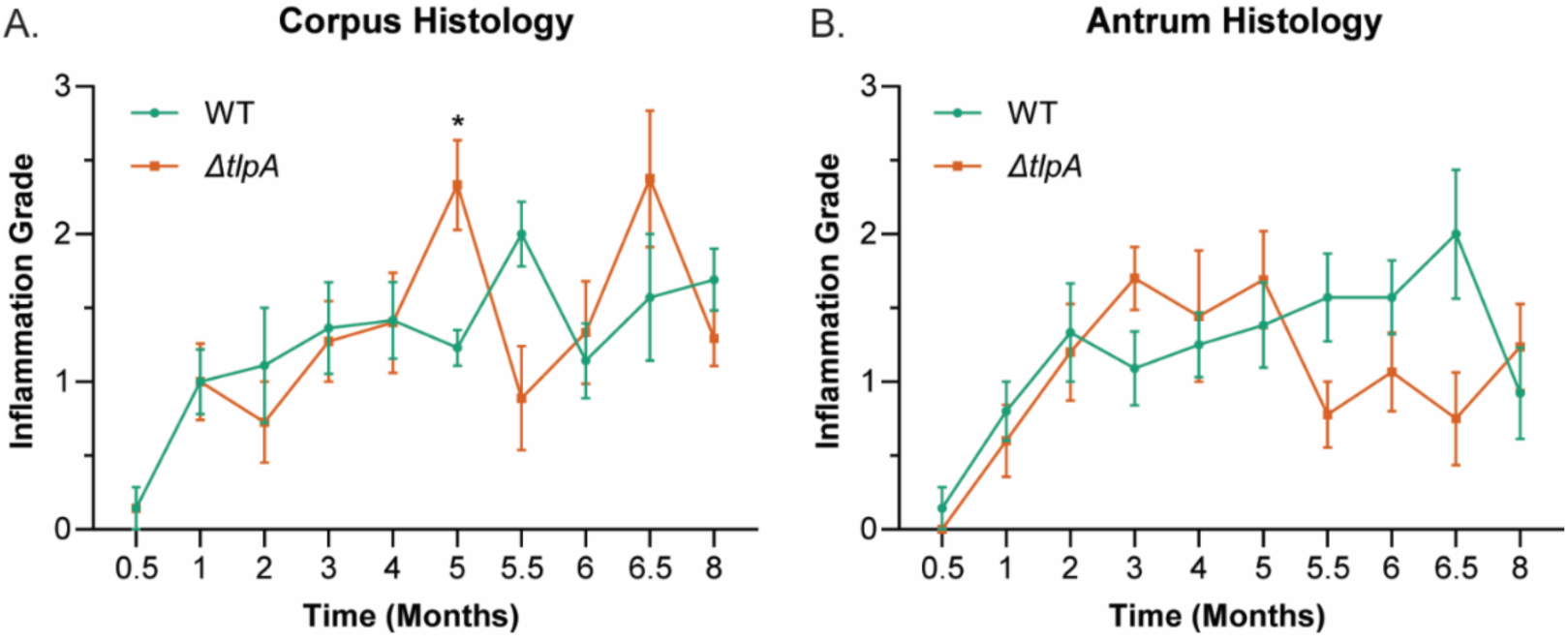
Δ*tlpA H. pylori* infections result in inflammation peaks with offset kinetics compared to WT. Female C57BL/6N mice (n ≥ 7 per group) were orally infected with WT SS1 GFP^+^ or Δ*tlpA* SS1 GFP^+^ for 0.5-, 1-, 2-, 3-, 4-, 5-, 5.5-, 6-, 6.5-, and 8-months, in two independent infections. Inflammation induced by WT SS1 GFP^+^ or Δ*tlpA* SS1 GFP^+^ infection groups was measured by assessing lymphocyte recruitment to the lamina propria of (A) corpus and (B) antrum via histology, as performed previously (33). Error bars represent the standard error of the mean. *, P<0.05, comparisons between strains at the same time point using two-way ANOVA, Šídák multiple comparison test.

Consistent with previous work, *tlpA* mutants caused heightened inflammation around 5-6 months post infection (Fig. 1A). Previous analysis did not dissect whether the *tlpA* inflammation was localized (33), and here we find that the elevated inflammation was restricted to the corpus, and did not occur in the antrum (Fig. 1). Surprisingly, the *tlpA* inflammation showed periodic variations, with high inflammation points followed by low ones (Fig. 1A). These fluctuations were most prominent in the corpus, as compared with the antrum (Fig. 1). When this inflammation profile was compared to that of WT, we found that WT infections also displayed inflammation periodicity in the corpus, but not antrum (Fig. 1). Indeed, the elevated inflammation of *tlpA* mutants appeared due to a rapid rise inflammation after 4 months of infection. WT had a similar response, but at a slightly later time. This offset resulted in the *tlpA* mutant inflammation exceeding WT at some times, e.g. 5 and 6.5 months. Due to the fluctuating nature, however, WT inflammation was greater than that of the *tlpA* mutant at other times, e.g. 5.5 months. The *tlpA* mutant corpus inflammation showed a greater range than that of WT in the period 4 months post infection. The *tlpA* mutant scores ranged from 0.9-2.3, while WT ranged only from 1.5-2.0. The antrum inflammation was different. In this case, both WT and *tlpA* mutant had similar increasing trends from 0.5-5 months. After that, WT mostly continued to increase slightly, while the *tlpA* mutant declined between 5-5.5 months, and then remained at this new lower level. Overall, these results suggest that the corpus and antrum have somewhat different inflammation processes, with the corpus displaying periodic variations of high and low inflammation grades. The *tlpA* mutant causes elevated inflammation, with top scores that are higher than those of WT, but also offset fluctuation kinetics.

### Increased Th17 cell density is observed in Δ*tlpA H. pylori* infected corpus during late infection

We next sought to better understand the nature of the periodic corpus immune responses, using flow cytometry to assess innate immune and effector T-cell populations in the samples from time course 1 (Supplemental Fig. 1) . On the innate immune side, we examined three types of DCs (CD103^+^ CD11b^+^ DCs, CD103^+^ CD11b^-^ DCs, CD103^-^ CD11b^+^ DCs), two types of monocytes (MHCII^-^ monocytes, MHCII^+^ monocytes), MHCII^+^ macrophages, neutrophils, and eosinophils, as described in previous work (39–41) (Sup. Fig. 2A). All of these innate immune populations were elevated relative to uninfected animals (Sup. Fig. 3), but mostly did not show patterns of temporal variation or substantial differences between WT and Δ*tlpA H. pylori* infection groups (Fig. 2). An exception was eosinophils, which were lower in WT infections at 6- and 8-months post-infection (Fig. 2A), although there was not elevated WT at these timepoints (Fig. 1). These results suggest the inflammation score fluctuation does not correlate with differences in the abundance of innate immune cell populations.

**Figure 2.**
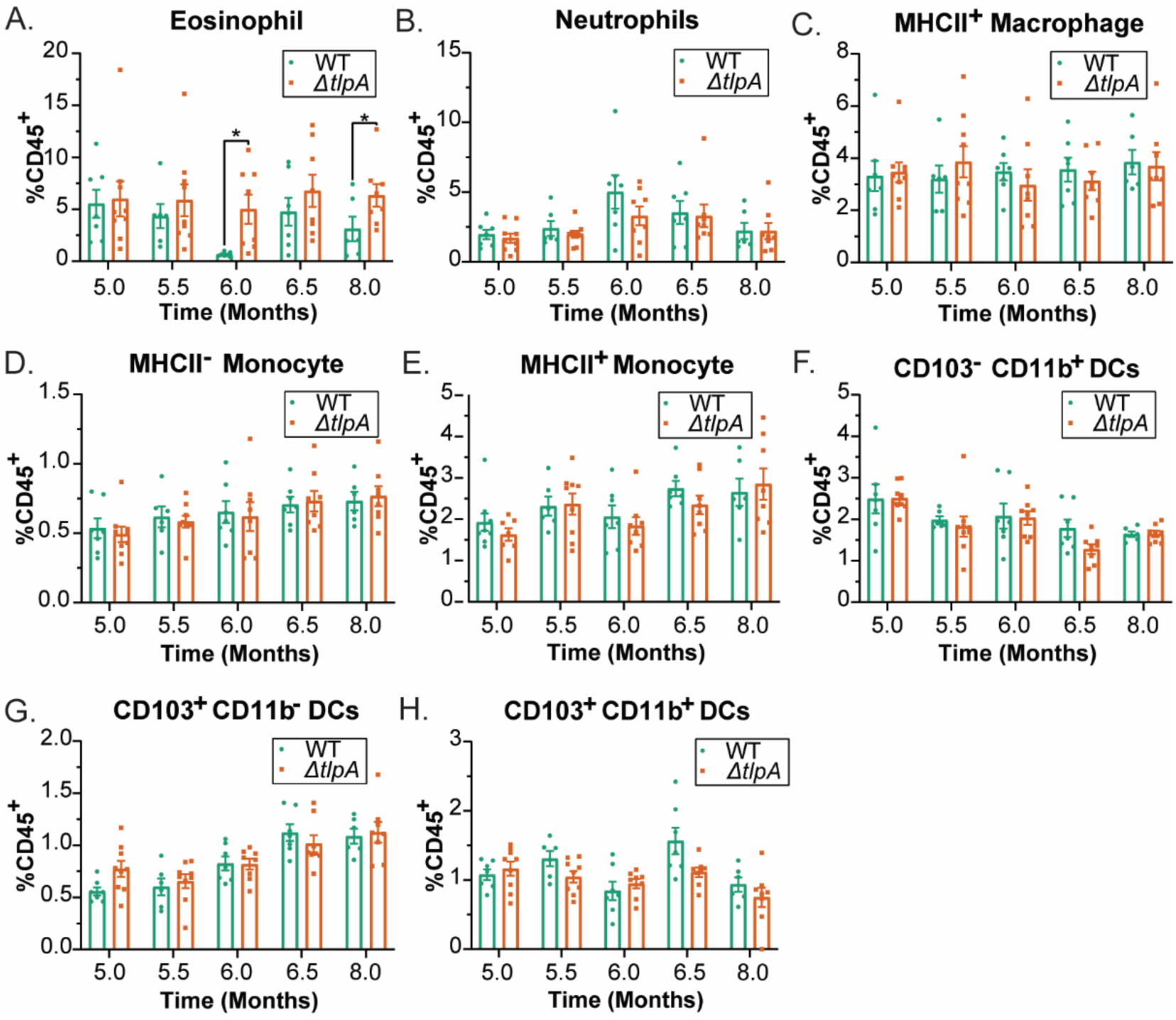
Innate immune cells recruitment during chronic infection is similar between WT SS1 and Δ*tlpA H. pylori* infected mice. Mice infected with WT SS1 GFP^+^ or Δ*tlpA* SS1 GFP^+^ were used to isolate corpus lamina propria leukocytes after 5-, 5.5-, 6-, 6.5-, and 8-months post-infection from the same mice as shown in Sup. Fig. 1A (n = 7-9 per group from one experiment). Analysis including uninfected mice is included in Sup. Fig. 2. A) neutrophils, B) eosinophils, C) MCHII^+^ macrophages, D) MCHII^-^ monocytes, E) MHCII^+^ monocytes, F) CD103^-^ CD11b^+^ DCs, G) CD103^+^ CD11b^-^ DCs, and H) CD103^+^ CD11b^+^ DCs were analyzed by flow cytometry. Data are presented as the frequency of each cell type as the percent of all live CD45^+^ lymphocytes. Error bars represent the standard error of the mean. *, P<0.05, comparisons between strains using two-way ANOVA, Šídák multiple comparison test.

We next examined the CD4^+^ effector T-cell populations, which are known to be key drivers of *H. pylori* inflammation (36, 43–46). Total CD4^+^ effector T-cells (CD4^+^ TCRβ^+^) were assessed, as well as each subset by the measurement of specific cell markers: Th1 (Tbet^+^), Th17 (RorγT^+^), and Treg (FoxP3^+^) (Sup. Fig. 2B). These cell types were all elevated in mice infected with either WT or *tlpA* mutant *H. pylori*, as compared to uninfected, with numbers steadily increasing by 10-fold over the time course (Sup. Fig. 4). At the 5 month time point, there were more Th17 cells than other types, but by 8 months, all three T cell subsets were found in equal amounts (Fig. 3, Sup. Fig. 4). There was some evidence of periodic fluctuation in cell amounts. At 5 months and 8 months post-infection, Th17 cells were significantly elevated in the Δ*tlpA H. pylori* infected mice as compared to WT (Fig. 3B). Th1 cell numbers also showed some periodicity, with peaks at 6 and 8 months. Treg numbers, in contrast, did not show variation, and were consistently equivalent between WT and mutant (Fig. 3A and C). Overall, these results suggest that some T cells may undergo periodic fluctuation, possibly accounting for the heightened inflammation grade of *tlpA* mutants at 5.5 months. There are not, however, dramatic changes in T cell counts between time points and strains.

**Figure 3.**
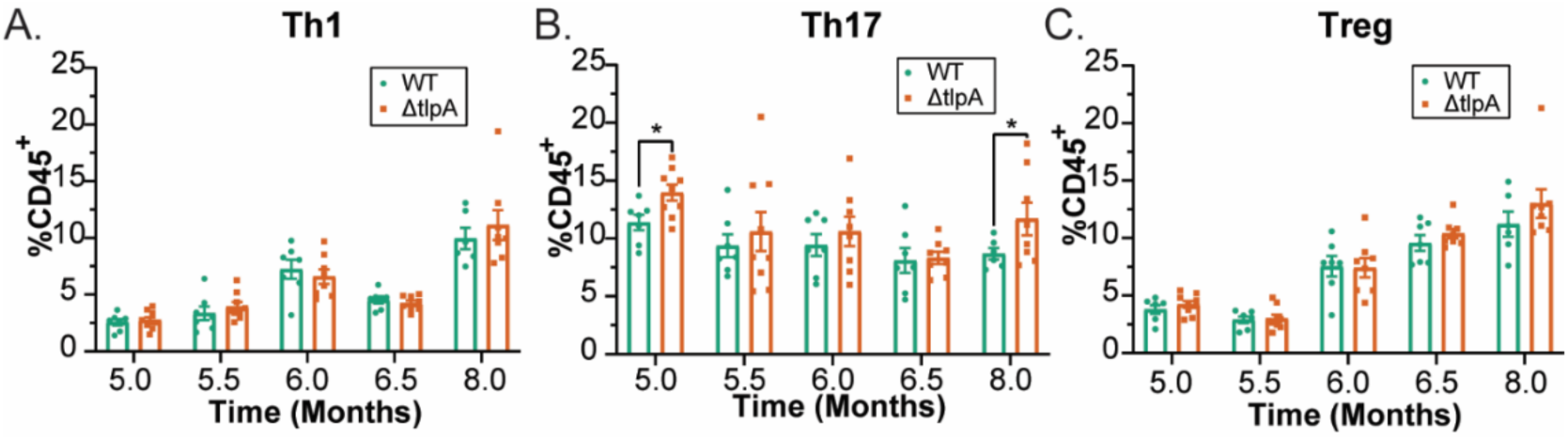
Δ*tlpA H. pylori* infected mice display increased Th17 cell populations recruited to the corpus lamina propria. Mice infected with WT SS1 GFP^+^ or Δ*tlpA* SS1 GFP^+^, and lymphocytes from the corpus lamina propria were isolated at 5-, 5.5-, 6-, 6.5-, and 8-months post-infection from the same mice from Fig. 2 (n = 7-9 per group, from one experiment). Analysis including uninfected mice is included in Sup. Fig. 3. A) Th1 (Tbet^+^ TCRβ^+^ CD4^+^), B) Th17 (RorγT^+^ TCRβ^+^ CD4^+^), and C) Tregs (FoxP3^+^ TCRβ^+^ CD4^+^) were analyzed by flow cytometry. Data are presented as the frequency of each cell type as the percent of all live CD45^+^ lymphocytes. Error bars represent the standard error of the mean. *, P<0.05, comparisons between strains using two-way ANOVA, Šídák multiple comparison test.

### Gland colonization patterns of *H. pylori* are dynamic throughout a long-term infection

Our analysis in Figure 1 shows that *H. pylori* corpus inflammation undergoes periodic fluctuations during late infection stages, and these fluctuations are location specific. Because inflammation is driven by *H. pylori* presence, we next analyzed *H. pylori* colonization levels and patterns. Previous work had assessed the total tissue colonization of both WT and *tlpA* mutant strains (33–35), but since that initial analysis, there has been increased appreciation that *H. pylori* is found in multiple microniches including within and outside of gastric glands, and between the corpus and antrum (8, 28, 32). Accordingly, we assessed the colonization patterns of WT and Δ*tlpA* SS1 GFP^+^ *H. pylori* from the time course samples above (Fig. 1). At each time point, we enumerated corpus and antrum total colony forming units (CFU) from bulk tissue, and then multiple gland parameters including the average number of bacteria per glands (gland colonization), the average number of bacteria per occupied gland (gland density), and the percentage of glands occupied (gland percent) (Fig. 4). Of note, gland colonization is the composite of both gland density and gland percent, thus providing an overall picture of gland colonization dynamics. The latter two metrics give contextual understanding to the overall gland colonization patterns.

**Figure 4.**
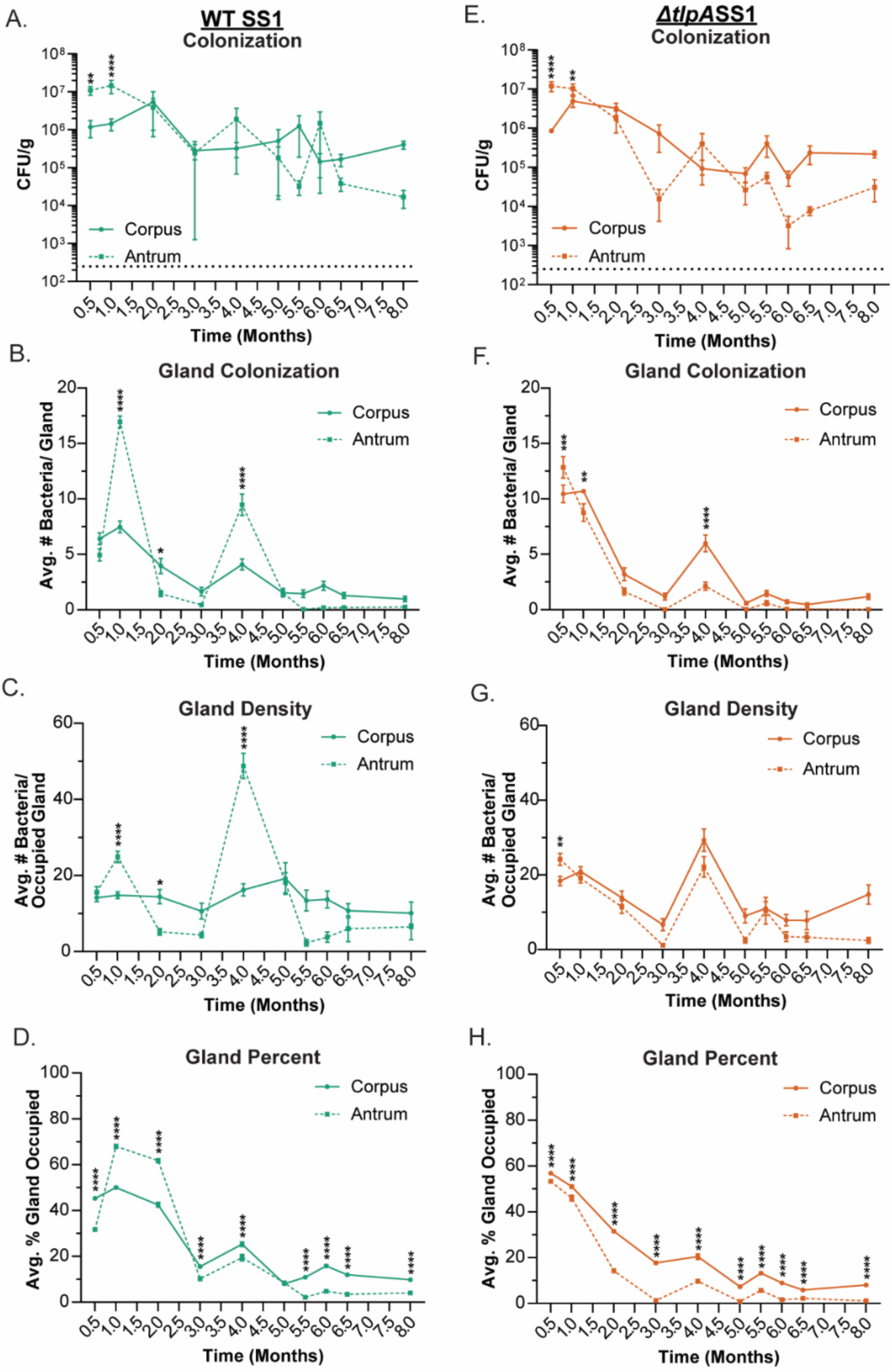
WT SS1 and Δ*tlpA H. pylori* SS1 have defined colonization patterns in the murine stomach. Colonization dynamics of WT SS1 and Δ*tlpA* SS1 within the mouse stomach were analyzed over a time course from the acute to chronic stage of infection from the same mice from Sup. Fig. 1A-B. Female C57BL/6N mice (n ≥ 4) were orally infected with WT SS1 GFP^+^ or Δ*tlpA* SS1 GFP^+^ for 0.5-, 1-, 2-, 3-, 4-, 5-, 5.5-, 6-, 6.5-, and 8-months. Total colonization levels of (A) WT and (E) Δ*tlpA* in the corpus and antrum region of the stomach was determined by homogenizing tissue and plating for colony forming units (CFU). For each strain, gland colonization phenotypes were quantified and compared between the corpus and antrum. Individual glands were isolated and GFP^+^ bacteria were visualized using the Bacterial Localization in Isolated Glands (BLIG) method (32). For each mouse, 100 isolated glands from the corpus and 100 from the antrum were analyzed. (B and F) Gland colonization is the average number of WT SS1 GFP^+^ or Δ*tlpA* SS1 GFP^+^ per all gland**s** analyzed in either region. (C and G) Gland density is the average number of WT SS1 GFP^+^ or Δ*tlpA* SS1 GFP^+^ per occupied gland. (D and H) Gland percent is the average percent of glands that are occupied by either WT SS1 GFP^+^ or Δ*tlpA* SS1 GFP^+^ at each time point. For all graphs, error bars represent the standard error of the mean. *, P < 0.05; **, P < 0.01; ***, P < 0.001; ****, P < 0.0001, comparisons between corpus and antrum using two-way ANOVA, Šídák multiple comparison test.

As reported previously, WT *H. pylori* initially colonizes the antrum to a higher level than in the corpus during the first ∼ 1 month of infection, and then antral numbers decline such that there are not significant differences in the total number of bacteria between these regions (Fig. 4A) (32, 48, 50). Δ*tlpA* mutants displayed a similar pattern (Fig. 4E), with two notable differences. First, the corpus CFU were elevated at 1 month post infection compared to WT (Fig. 5A), and the early decline in antral numbers was more dramatic (Fig. 5E). Overall, these bulk tissue analyses support previous findings that the *tlpA* mutant does not have substantial colonization defects, and indeed may even colonize to slightly higher levels in the corpus in early infections.

**Figure 5.**
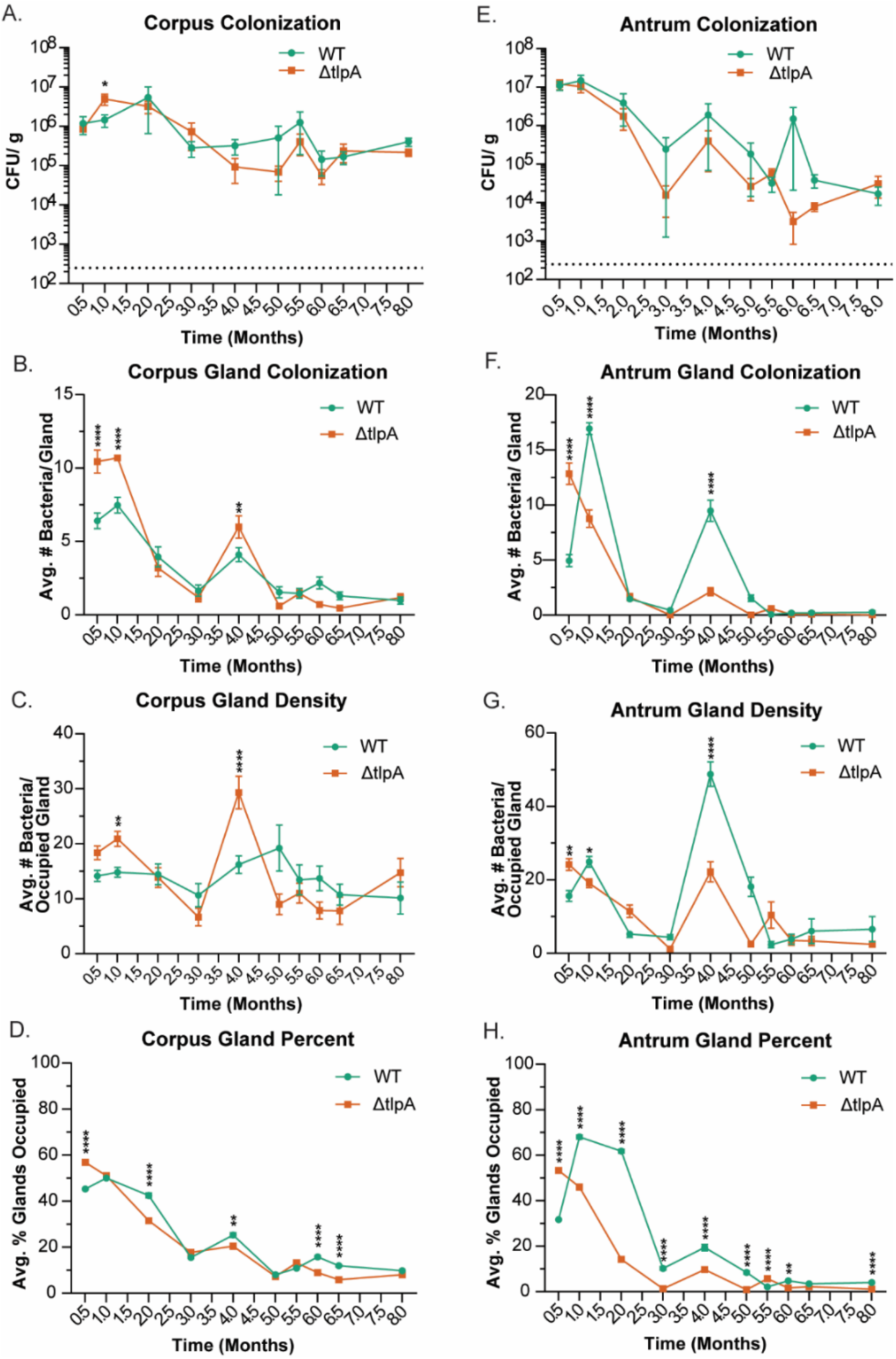
Δ*tlpA H. pylori* occupy more glands, with greater density during early infection, and colonize glands with lower frequency during late infection. Overall colonization and gland colonization phenotypes were quantified and compared between WT SS1 GFP^+^ or Δ*tlpA* SS1 GFP^+^ from the infection groups from the same mice as in Fig. 4. Colonization of WT SS1 and Δ*tlpA* in the corpus was assessed for (A) total colonization, (B) gland colonization, (C) gland density, and (D) gland percent. Colonization of WT SS1 and Δ*tlpA* in the antrum was assessed for (E) total colonization, (F) gland colonization, (G) gland density, and (H) gland percent. For all graphs, error bars represent the standard error of the mean. *, P<0.05; **, P<0.01; ***, P<0.001; ****, P<0.0001, comparisons between strains using two-way ANOVA, Šídák multiple comparison test.

We next analyzed the gland populations. Similar to previous reports, WT colonized ∼30-50% of the glands by two weeks (Fig. 4D), with ∼ 20 bacteria per gland in both the corpus and antrum (Fig. 4B, 4C) (32). Antral gland colonization expanded during the next two weeks (Fig 4B). The percent of glands occupied increased to ∼70%, and the number of bacteria per occupied gland increased to ∼30 (Fig 4C-D). In contrast, the corpus gland populations remained steady (Fig. 4B-D). After this expansion period, antral gland colonization showed an abrupt decline between months 1-3 (Fig. 4B). This decline was mostly due to a significant decline in the number of bacteria per gland, while the percent of occupied glands remained high (Fig. 4C-D). The corpus, in contrast, retained a steady number of bacteria per occupied gland at ∼20, and declined only in the percent of glands occupied down to ∼15%. During this same period, we observed multiple differences between WT and *tlpA* mutant. The most noticeable difference was that the *tlpA* mutant colonized the corpus glands to higher levels than did WT in early infection. In the first month, Δ*tlpA H. pylori* showed elevated numbers associated with high bacterial numbers per gland and sometimes high percent of occupied glands (Fig. 5C and D).

From 2- 8-months post-infection, there were few differences in gland colonization between the strains in either the corpus or antrum (Fig. 5B and F). For both strains, gland colonization decreased from the peak at 1-month post-infection over the remainder of the infection (Fig. 5B and F). There was an increase in both WT and Δ*tlpA H. pylori* gland colonization at 4 months in one time course (Fig. 4) but not the other (Supplemental Fig. 4). Δ*tlpA H. pylori* consistently colonized fewer glands than WT in both regions, achieving significance in many instances (Fig. 5D and H). One exception to this pattern was at 5.5-months post-infection in the antrum when Δ*tlpA H. pylori* had significantly higher gland percent occupancy (Fig. 5H), and a non-significant increase in gland density (Fig. 5G).

Overall, these results demonstrate that Δ*tlpA H. pylori* gland colonization pattern has two main differences from that of WT. First, at early times, the Δ*tlpA* mutant has higher gland colonization than WT in both the corpus and the antrum (Fig. 5B, 5F). The corpus gland colonization was higher at 0.5 and 1 months, while the antrum was only higher at 0.5 months. In later infection, however, the Δ*tlpA* mutant gland colonization was generally below that of WT. In some infections, there was a spike in numbers at 4 months but this was not a consistent finding (Fig. 5, Suppl. Fig. 5).

## Discussion

In this work, we sought to understand how *H. pylori* influences inflammation. Our study built on an earlier observation that Δ*tlpA* mutants have elevated late-infection inflammation (33). Here we report that inflammation induced by Δ*tlpA* mutants is more complex than previously appreciated. Interestingly, we found that *tlpA* mutants and WT display inflammation that fluctuates, with the *tlpA* inflammation out of phase from that of WT. This out-of-phase variation gave rise to times when *tlpA* mutant inflammation was either higher or lower than WT. This outcome occurred most dominantly in the mouse corpus; the antrum, in contrast, displayed less variation, and similar inflammation levels between the WT and *tlpA* mutant.

We characterized the Δ*tlpA*-mutant immune response using flow cytometry. We observed Δ*tlpA H. pylori* infected mice had increased Th17 cell density (Fig. 3B), but not any other cell type, during the period of increased inflammation at 5-months post-infection (Fig. 2 and Fig. 3A+C). Th17 cells play a major role in inflammation development during *H. pylori* colonization, primarily through the production of pro-inflammatory cytokines (51–53). Some of the primary cytokines include: 1) IL-17A, which promotes neutrophil recruitments through IL-8 and promotes the expression of antimicrobial peptides from epithelial cells; 2) IL-17F, which upregulates the expression of other pro-inflammatory cytokines and chemokines, inducing IL-2, TGF-β, IL-6, and GM-CSF; 3) IL-21, which promotes the proliferation of Th1 and Th17 cells, creating a positive feedback loop, and promote the proliferation of B-cells; and 4) IL-22, which acts on non-hematopoietic cells by upregulating antimicrobial peptide expression and promoting tissue repair (51). Interestingly, recent work has shown that arginine, a TlpA chemoattractant (25), and the polyamine pathway are critical for maintaining Th17 pathogenicity (54). Both of these factors drive Th17 cells toward a pro-inflammatory state. One idea is that WT *H. pylori* uses TlpA to mediate a chemoattractant response towards arginine, allowing *H. pylori* to readily use this amino acid and thus limit its availability to the host. This usage, in turn, could reduce arginine uptake by Th17 cells, resulting in decreased cytokine production and pathogenicity (Wagner et al., 2021).

We also observed significantly higher eosinophil recruitment compared to WT at multiple time points in Δ*tlpA H. pylori* infected animals (Fig. 2A). Recently, eosinophils have been shown to play an important role during *H. pylori* colonization, as eosinophils interact with *H. pylori* in the gastric mucosa, regulating Th1 cell proliferation (39). Th1 cells are a significant contributor to the development of pathologies that are observed during *H. pylori* colonization through the production of IFNγ (45). However, changes in eosinophil populations do not correlate with any change in Th1 recruitment during the period of observation at the 5-8-month post-infection window (Fig. 2A and Fig. 3), suggesting a more dominant role for the Th17 cell population in *tlpA* mutant inflammation.

In this study, our latest time point was at 8-months post-infection. However, like in the human stomach, *H. pylori* can establish chronic colonization, lasting for over a year in a murine model. Colonization studies up to 15-month-post-infection with *H. pylori* SS1 have been performed, showing that mice colonized for 15-months had higher inflammation than at 6-month post-infection (55). Given this outcome, it would be interesting to see how inflammation evolves over a more extended period and whether periodic fluctuations continue.

Our data suggest that TlpA is important for causing consistent levels of inflammation in the stomach. Keeping inflammation at a low level is important for colonization maintenance. Following the inflammatory spike in the *tlpA* mutant at 5-months post-infection, we observed ∼7- and ∼17-fold colonization decreases in the corpus and antrum, respectively. It has been well established in the literature that inflammation severity and colonization levels are inversely related (36, 56, 57). Therefore, it is presumably advantageous for *H. pylori* to limit the amount of inflammation it induces during infection.

Over the past decade, it has become appreciated that *H. pylori* colonization of gastric glands is regulated by chemotaxis (8). Non-chemotactic mutants, including Δ*cheY* and Δ*chepep*. have severe defects in gland colonization compared to WT *H. pylori* during the first month of infection (8, 17, 32, 31, 47). These mutants are fully non-chemotactic, meaning they are unable to detect any receptor input. Mutants that lack specific chemoreceptors also have infection phenotypes, but are generally not as severe. Each chemoreceptor expressed by *H. pylori* is responsible for sensing multiple ligands and signals (19). Accordingly, progress has been made to understand how chemoreceptors regulate the localization of *H. pylori* in the stomach. Studies with the chemoreceptor TlpD are illustrative of the roles that individual chemoreceptors can play. Δ*tlpD H. pylori* have overall colonization defects in the stomach (35), associated with a decrease in the percent of glands colonized, but interestingly, higher bacterial numbers per gland. This colonization phenotype is ameliorated in mice defective in reactive oxygen species (ROS) production, consistent with TlpD’s reported role in sensing and initiating a chemorepellent response to ROS (26, 31). In addition to ROS, TlpD has been shown to sense pH synergistically with TlpA. Δ*tlpAD H. pylori* have a more severe overall colonization defect at 2-weeks post-infection than Δ*tlpD H. pylori* and are defective in gland colonization. These defects were improved via omeprazole treatment, a proton-pump inhibitor that increases the pH of the stomach (28). In total, this work demonstrates the importance chemoreceptors can play during colonization. The inability to sense specific signals in the gastric environment leads to altered localization, leaving *H. pylori* susceptible to non-optimal or even toxic host conditions and immune responses.

Our studies examined the colonization patterns of both WT and Δ*tlpA H. pylori* throughout a long-term infection. We found that WT and Δ*tlpA H. pylori* have similar overall colonization trends within the corpus and antrum. Both strains favor the antrum during the first month of colonization, but the *tlpA* mutant achieves higher gland colonization (Fig. 4A and E). After this point, there are no statistical differences in overall colonization between regions of the stomach when analyzing each strain independently. Δ*tlpA H. pylori* favors corpus glands throughout the infection time course, except at 0.5-months post-infection (Fig. 4F-G). WT, however, tends to favor antral glands 1-2-months post-infection, with either higher gland percent or density (Fig. 4B-D). Thus, it seems that loss of *tlpA* results in strains that are less able to thrive in the antrum and more able to thrive in the corpus. It is possible that TlpA’s signals—arginine, cysteine and fumarate (25)— are critical in the antral glands and therefore *tlpA* mutants cannot access these ligands efficiently and do not grow as well as WT.

Intriguing to note, TlpA appears to be under selective pressure, accumulating a high frequency of mutations in its dCache sensing domain (58). While it is not yet known how these mutations affect TlpA sensing and signaling, it is interesting that *tlpA* is one of the more highly affected loci and may be subject to mutations that change its properties and possible inflammatory-control abilities.

There is accumulating evidence that *H. pylori* gland colonization is correlated with inflammation. Non-chemotactic *H. pylori* mutants colonize the glands poorly within the first month, and induce low inflammation levels at 2-month post-infection, which is characterized by a decreased Th17 response (46) (32). In contrast, our results show that Δ*tlpA H. pylori* have high gland colonization in the corpus until 1-month post-infection, characterized by increased gland density and gland percent relative to WT (Fig. 5B-D). A potential mechanism by which chemotaxis affects inflammatory outcomes is through controlling colonization of gastric glands during the acute stage of infection. During this period, various myeloid cell types in the stomach interact with *H. pylori*. This is critical for developing the effector T-cell response, characterized by the recruitment and proliferation of the pro-inflammatory, Th1 and Th17 cells, and the anti-inflammatory Tregs during infection (18). It is plausible that situations that result in high numbers of bacteria per gland increases the chance for total or specific myeloid cells to interact with *H. pylori.* High or distinct myeloid sampling might, in turn, result in differences in T-cell priming and subsequent T-cell populations. On the other hand, non-chemotactic mutants have low gland density and occupancy compared to WT (32). As a result, there may be decreased T-cell priming during the early stage of infection, potentially explaining why non-chemotactic *H. pylori* induce a dampened Th17 response during infection (46).

## Methods

### Bacterial strains and culture conditions

*H. pylori* strains used in this study were all derived from the *H. pylori* strain SS1. They include WT, WT GFP^+^, Δ*tlpA* GFP^+^ (This study), and Δ*tlpA::kan-sac* (59, 32, 60). Bacteria were grown at 37°C under microaerobic conditions of 5% O_2_, 10% CO_2_, and 85% N_2_ on solid media containing Columbia Blood Agar Base (BD), containing 5% defibrinated horse blood (Hemostat Laboratories, Dixon, CA), 50 µg/ml cycloheximide (VWR), 10 µg/ml vancomycin, 5 µg/ml cefsulodin, 2.5 U/ml polymyxin B (all from Gold Biotechnology), and 0.2% (wt/vol) beta-cyclodextrin (Spectrum Labs, Gardena, CA) (CHBA) or in Brucella broth (BD BBL/Fisher) with 10% heat-inactivated fetal bovine serum (FBS) (Life Technologies) (BB10), with shaking. Chloramphenicol-resistant mutants were selected using 10 μg/ml chloramphenicol (Gold Biotechnology) on CHBA, as previously described (34).

The Δ*tlpA* SS1 clean deletion mutant was made by natural transformation of Δ*tlpA::kan-sac H. pylori* SS1 with pTA10, which contains a *tlpA* deletion flanked by 500 bp of the up- and down-stream regions of *tlpA* (60). Sucrose resistant colonies were selected on CHBA containing 75 µg/ml sucrose (Fisher Chemical), followed by screening for kanamycin sensitivity on CHBA with 15 µg/ml kanamycin (Gold Biotechnology). The *ΔtlpA* deletion was confirmed via PCR (TlpA_SS1_5’, TTGTCTAAAGGTTTGAGTATC; TlpA_SS1_3’, TTAAAACTGCTTTTTATTCAC) (Data not shown) and western blot using anti-TlpA22 (33) (Sup. Fig. 6). Δ*tlpA* SS1 was then transformed with the GFP expression plasmid, pTM115 (*ureA_p_-*GFP *aphA3* Km^R^) (32). Colonies were screened for kanamycin resistance and GFP^+^ fluorescence

### Ethics statement

The University of California Santa Cruz Institutional Animal Care and Use Committee approved all animal protocols and experiments performed (Protocol Ottek1804). Female C57BL/6N mice (*Helicobacter* free, Charles River) were housed and cared for at the University of California Santa Cruz Vivarium.

### Animal infections

Female C57BL/6N mice (*Helicobacter* free, Charles River) were orally infected with *H. pylori* at 6 to 8 weeks of age, as done previously (8, 31). *H. pylori* SS1 strains were grown to late exponential phase (OD_600_ = ∼0.75) in BB10 and were checked for GFP fluorescence and motility using a Nikon Eclipse E600 phase-contrast microscope at 400x magnification with a LED illuminator (pE-300^white^, CoolLED) with a fluorescent filter for GFP. Motile GFP^+^ cultures were concentrated to OD_600_ = 3 (∼9 x 10^8^ CFU/ml) by centrifugation of 1 ml culture aliquots in 1.5 ml Eppendorf Tubes at 2320 x g for 10 minutes. Culture supernatant was removed via aspiration and pellets were gently resuspended in BB10 to achieve an OD_600_ = 3. For infection, mice were scruffed and oriented at 45° with head and abdomen up, then orally fed 50μl of the OD_600_ = 3 culture via a pipet tip, for an inoculum of 4.5 x 10^7^ CFU. Mice in the uninfected group were fed 50μl of BB10. Input inocula were plated on CHBA plates to determine the true input CFU. Two independent time course infections were performed for this study. The infection time course reported in Figures 1-5 included time points at 0.5- (WT and Δ*tlpA*, n = 7), 1- (WT and Δ*tlpA*, n = 7), 2- (WT and Δ*tlpA*, n = 4), 3- (WT and Δ*tlpA*, n=4), 4- (WT and Δ*tlpA*, n=7), 5- (WT, n=7; Δ*tlpA*, n=9), 5.5- (WT, n=7; Δ*tlpA*, n=9), 6- (WT, n=7; Δ*tlpA*, n=8), 6.5- (WT, n=7; Δ*tlpA*, n=8), and 8-months (WT and Δ*tlpA,* n=6). The infection time course reported in Figure 1 and 6. and Supplemental Figures 5-6, include, 2- (WT and Δ*tlpA*, n=7), 3- (WT and Δ*tlpA* n=7), 4- (WT and Δ*tlpA*, n=6), 5- (WT and Δ*tlpA*, n=6), 6- (WT and Δ*tlpA*, n=7), and 8-months (WT, n=7; Δ*tlpA*, n=9). From this time course, only 3 mice were used for gland analysis from 2-5-months post-infection, after which all mice from the infection group were used. For both time course infections, 3-4 uninfected mice were included per time point. After the infection period, the mice were sacrificed by CO_2_ narcosis, the stomach was removed at the stomach-esophageal junction and the antrum-duodenum sphincter, then opened by cutting along the lesser curvature of the stomach. Stomach contents were washed gently using ice-cold PBS. For dissection, a piece of tissue from the antrum to corpus was removed and stored in a histology cassette (Histoware) with sponge pads in Buffered Formalde-Fresh solution (Fisher Chemical) for histology. The remaining stomach was then separated between the antrum and corpus at the transition zone, based on tissue coloration. Each section was then divided into pieces to determine output CFU, to assess gland colonization, and for flow cytometry. To determine colonization load, tissue pieces were weighed, homogenized using the Bullet Blender (Next Advance) with 1.0 mm zirconium silicate beads, and then plated onto CHBA plates, supplemented with 20 µg/ml bacitracin, 10 µg/ml nalidixic acid, and 15 µg/ml kanamycin, to determine CFU/g of stomach tissue.

### Gland isolation and microscopy

Gastric glands from the antrum and corpus were isolated and *H. pylori* colonization within glands was quantified using the Bacterial Localization in Isolated Glands methods, as done previously (31, 32). Briefly, glands were isolated by agitating dissected tissue from the antrum or corpus in PBS with 5 mM EDTA at 4°C for 2 hours. The incubated tissue was then transferred to PBS with 1% sucrose and 1.5% sorbitol and vigorously by hand for 30 sec (both Fisher Chemical). Glands were labeled with 10 µg/ml Hoechst DNA stain (Life Technologies) and stored on ice. Glands were visualized with a Nikon Eclipse E600 microscope with a LED illuminator (pE-300^white^, CoolLED) and fluorescent filters for 4’,6’-diamido-2-phenylindole (DAPI) and GFP. At each infection time point, 100 glands from the antrum and 100 from the corpus were imaged and the number of GFP^+^ *H. pylori* per gland were manually counted. Gland colonization was calculated as the average number of bacteria per gland. Gland density was calculated by averaging the number of bacteria within occupied glands. Gland percent is calculated as the average frequency of glands occupied per mouse. Standard error of the mean was calculated for each. Statistical analysis of the data for between infection group or stomach regions at each time point was performed using two-way analysis of variance (ANOVA), Šídák multiple comparison test.

### Histology

Tissue pieces from the antrum to corpus stored in Buffered Formalde-Fresh solution (Fisher Chemical) in histology cassettes with sponges were processed for sectioning and graded for lymphatic infiltration as done previously (33). Briefly, tissue was embedded in paraffin, sectioned (5 μm), stained in hematoxylin and eosin, and evaluated by a pathologist in a blind fashion (J. Elliot Carter). To ensure reproducibility, slides were read twice and checked to ensure identical grades were obtained. Inflammation, defined as lymphocytic infiltration, was assessed using standard methods (33, 61) and sections were given scores as follows: 0, no infiltrate; 1, mild, multifocal infiltration; 2, mild, widespread infiltration; 3, mild, widespread and moderate, multifocal infiltration; 4, moderate, widespread infiltration; and 5, moderate, widespread and severe, multifocal infiltration. Neutrophil infiltration was scored as present or absent. Sections were also assessed for atrophy according to previously defined standards (62). Rugge *et al.* defined gastric atrophy as the loss of gastric glands in the area of the gastric mucosa being sampled. Atrophy can be associated with metaplasia, where gastric glands from a section of stomach are lost and replaced by gastric glands from a separate region of the gastric mucosa, or as atrophy without metaplasia, in which gastric glands from a section of stomach are lost with no replacement by other gastric glands and fibrosis of the lamina propria is seen. Samples negative for atrophy lack either manifestation of atrophy described. Mice from both time course infections were pooled for analysis. The average inflammation grade and standard error of the mean was calculated for each strain per time point. Statistical analysis of the data for between strains was performed using two-way ANOVA, Šídák multiple comparison test.

### Gastric lamina propria leukocyte isolation

Lamina propria leukocyte isolation from gastric tissue was performed as done previously (40, 39, 63). Tissue pieces from the antrum and corpus were placed in 50 ml conical tubes in 5 ml RPMI (Gibco) with 10% FBS (Gibco) on ice during the dissection of the mice. Media was discarded by aspiration and replaced by 30 ml of 1x Hanks Balance Salt Solution (Gibco) with 0.5% BSA (MillaporeSigma) and 5 mM EDTA (Fisher Chemical) prewarmed to 37°C. The tubes were then incubated horizontally at 37°C with shaking for 1 hour at 200 rpm in a Thermo Scientific MaxQ 4000 benchtop orbital shaker. After incubation, the supernatant was removed by aspiration, the tissue resuspended in 30 ml of room temperature PBS for 2 minutes, and then the PBS was replaced with 15 ml of RPMI with 10% FCS, 500 U/ml Type IV Collagenase (Sigma-Aldrich), 0.05 mg/ml DNaseI (Gold Biotechnology) pre-warmed to 37°C and incubated horizontally at 37°C with shaking for 1 hr at 200 rpm in a Thermo Scientific MaxQ 4000 benchtop orbital shaker. Following incubation, the solution was pulled through a 20 ml syringe 10-15 times to finish homogenization. The supernatant containing lamina propria leukocytes was filtered through a 40 µm cell strainer (Falcon) and then 30 ml of PBS was added. Cells were collected by centrifugation in a Thermo Scientific Sorvall Legend XTR with a TX-1000 Rotor at 1500 rpm (526 x g) for 8 min, the supernatant was discarded by aspiration, the cell pellet was resuspended in 4 ml of 80% Percoll (GE Healthcare), added to a 15 ml falcon tube, then overlayed with 4 ml of 40% Percoll. The 40/80% Percoll gradient was centrifuged for 15 min at 3000 rpm at 18°C. The interphase, containing cells, was collected and washed with PBS with 0.5% BSA.

### Flow cytometry

Flow cytometry was carried out as described previously (40, 39, 63). For surface staining, cells were stained in PBS with 0.5% BSA with Fixable Viability Dye eFluor 780 (eBioscience) and a combination of the following antibodies: anti-mouse CD45 (PE/Cy5, 30-F11, BioLegend), CD45 (Alexa Fluor 700, 30-F11, BioLegend), CD11c (APC, N418, BioLegend), I-A/I-E (Alexa Fluor 700, M5/114.15.2, BioLegend), F4/80 (FITC, BM8, BioLegend), CD103 (Brilliant Violet 605, 2E7, BioLegend), CD11b (Brilliant Violet 421, M1/70, BioLegend), Ly-6G (PE/Dazzle 594, 1A8, BioLegend), Ly-6C (PE/Cy7, HK1.4, Biolegend), Siglec-F (PE, S17007C, BioLegend), TCRβ (PE/Cy7, H57-597, BD), or CD4 (APC, RM4-5, BioLegend). Mouse Fc Block (CD16/CD32, BD) to minimize unspecific antibody binding. For transcription factor staining, cells were fixed and permeabilized with FoxP3 Fix/Perm Buffer Set (BioLegend) following the manufacturer’s instructions. Cells were stained with anti-mouse T-Bet (FITC, 4B10, BioLegend), RORγT (PE, Q31-378, BD), and FoxP3 (Pacific Blue, BioLegend, MF-14). Absolute cell counts were determined by adding CountBright Plus Absolute Counting Beads (Invitrogen) to each sample, which were analyzed on an LSRII (BD Biosciences). Up to 300,000 cells were recorded per mouse, with some samples containing as few as 30,000 cells. Absolute counts are reported at cells/ ul. Analysis was performed using FlowJo (BD).

## Acknowledgments

We thank Martha Zuniga, Vicki Auerbuch Stone, and Fitnat Yildiz for their thoughtful comments on this manuscript. We thank Bari Holm Nazario and the UCSC Institute for the Biology of Stem Cells Flow Cytometry Facility for technical support and training (RRID: SCR_021149). The described project was supported by the National Institute of Allergy and Infectious Disease (NIAID) grant RO1AI116946 to KMO. The funders had no role in study design, data collection, and interpretation, or decision to submit the work for publication.

## Supplemental Material

**Supplemental Figure 1.**
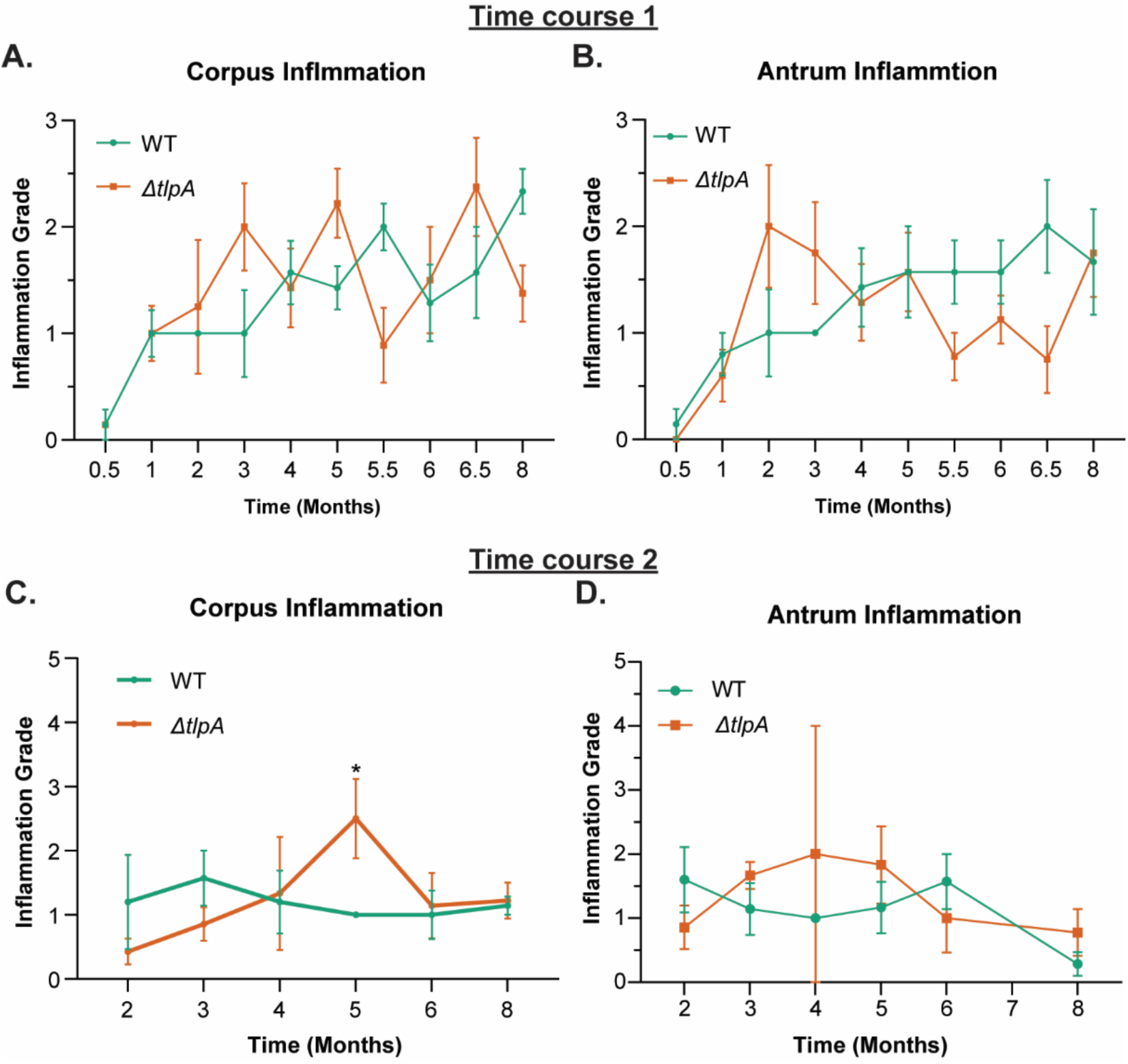
Significantly more inflammation is induced by Δ*tlpA H. pylori* in a transient manner during chronic infection. Two independent infection time courses were performed were female C57BL/6N mice orally infected with WT SS1 GFP^+^ or Δ*tlpA* SS1 GFP^+^. Time course 1 assessed time points at 0.5- (WT and Δ*tlpA*, n = 7), 1- (WT and Δ*tlpA*, n = 7), 2- (WT and *ΔtlpA*, n = 4), 3- (WT and Δ*tlpA*, n=4), 4- (WT and Δ*tlpA*, n=7), 5- (WT, n=7; Δ*tlpA*, n=9), 5.5- (WT, n=7; Δ*tlpA*, n=9), 6- (WT, n=7; Δ*tlpA*, n=8), 6.5- (WT, n=7; Δ*tlpA*, n=8), and 8-months (WT and Δ*tlpA,* n=6). Time course 2 assessed time points at 2- (WT and Δ*tlpA*), 3- (WT and Δ*tlpA*, n=7), 4- (WT and Δ*tlpA*, n=6), 5- (WT and Δ*tlpA*, n=6), 6- (WT and Δ*tlpA*, n=7), and 8- months (WT, n=7; Δ*tlpA*, n=9). Inflammation induced by WT SS1 GFP^+^ or Δ*tlpA* SS1 GFP^+^ infection groups was measured by assessing lymphocyte recruitment to the lamina propria of (A) corpus and (B) antrum via histology, as performed previously (33). Error bars represent the standard error of the mean. *, P<0.05, comparisons between strains using two-way ANOVA, Šídák multiple comparison test.

**Supplemental Figure 2.**
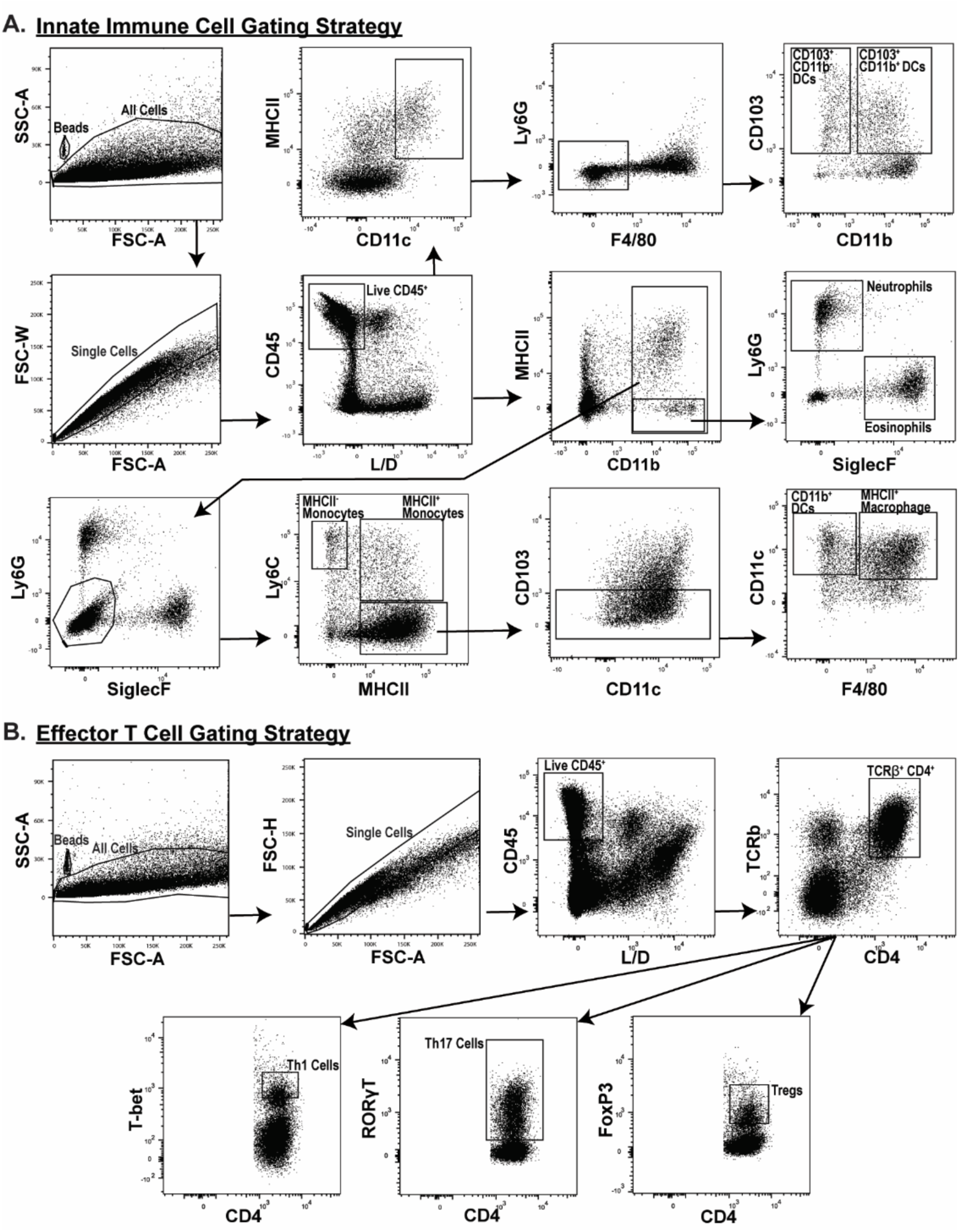
Gating strategy used to identify innate immune and effector T cell populations. Leukocytes were isolated from the corpus lamina propria via enzymatic digestion with collagenase IV and Percoll gradient centrifugation. Isolated leukocytes were then analyzed via flow cytometry. A) Gating strategy for innate immune cell types, as performed by Arnold et a. 2017. B) Gating strategy for CD4^+^ effector T cell populations.

**Supplemental Figure 3.**
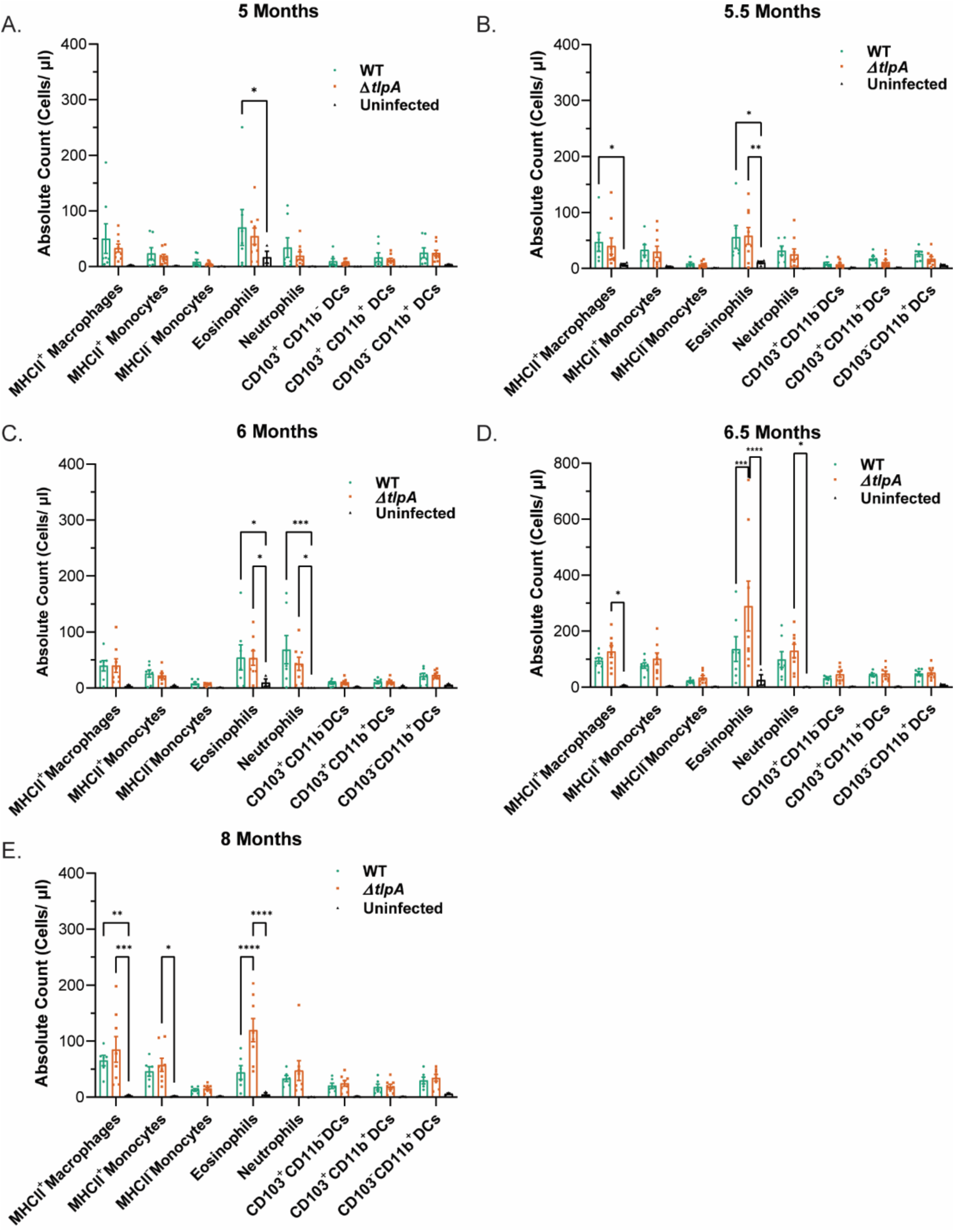
Absolute counts of innate immune cell populations recruited to the corpus during chronic infection. Mice were infected with WT SS1 GFP^+^ or Δ*tlpA* SS1 GFP^+^ and leukocytes from the corpus lamina propria were isolated at 5-, 5.5-, 6-, 6.5-, and 8-months post-infection (n = 7-9 per group). (A-E) neutrophils, eosinophils, MCHII^+^ macrophages, MCHII^-^ monocytes, MHCII^+^ monocytes, CD11b^+^ DCs, CD103^+^ CD11b^-^ DCs, and CD103^+^ CD11b^+^ DCs were analyzed by flow cytometry. Data are presented as the absolute count (cells/ µl) of each cell type. Error bars represent the standard error of the mean. *, P<0.05; **, P<0.01; ***, P<0.001; ****, P<0.0001, comparisons between strains using two-way ANOVA, Tukey multiple comparison test.

**Supplemental Figure 4.**
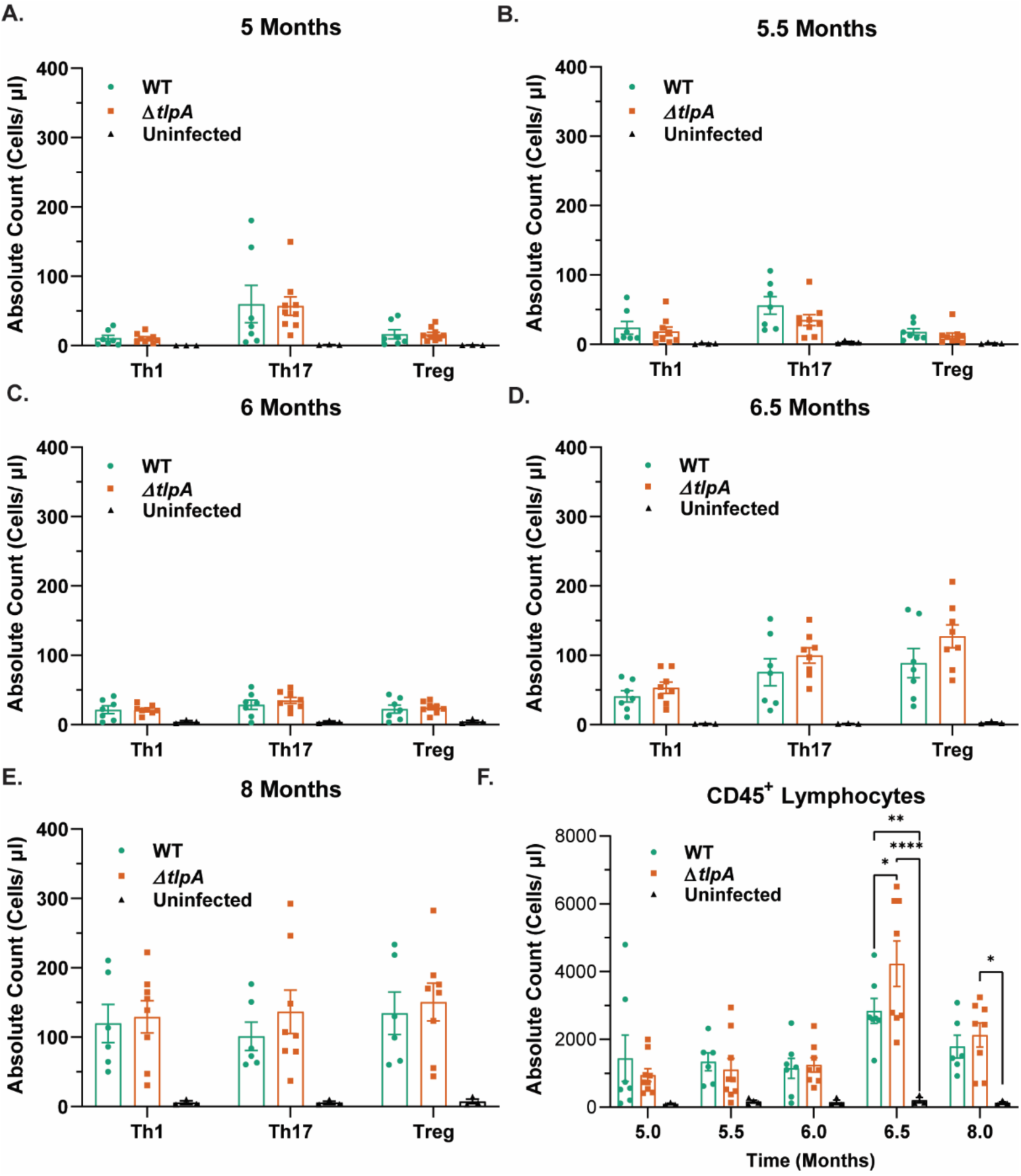
Absolute counts of effector T cell populations and total CD45^+^ leukocytes recruited to the stomach during chronic infection. Mice were infected with WT SS1 GFP^+^ or Δ*tlpA* SS1 GFP^+^. Leukocytes from the corpus lamina propria were isolated at 5-, 5.5-, 6-, 6.5-, and 8-months post-infection (n = 7-9 per group). (A-E) Th1, Th17, Tregs, and CD45^+^ leukocytes were analyzed by flow cytometry. Data are presented as the absolute count (cells/ µl) of each cell type. Error bars represent the standard error of the mean. *, P<0.05; **, P<0.01; ***, P<0.001; ****, P<0.0001, comparisons between strains using two-way ANOVA, Tukey multiple comparison test.

**Supplemental Figure 5.**
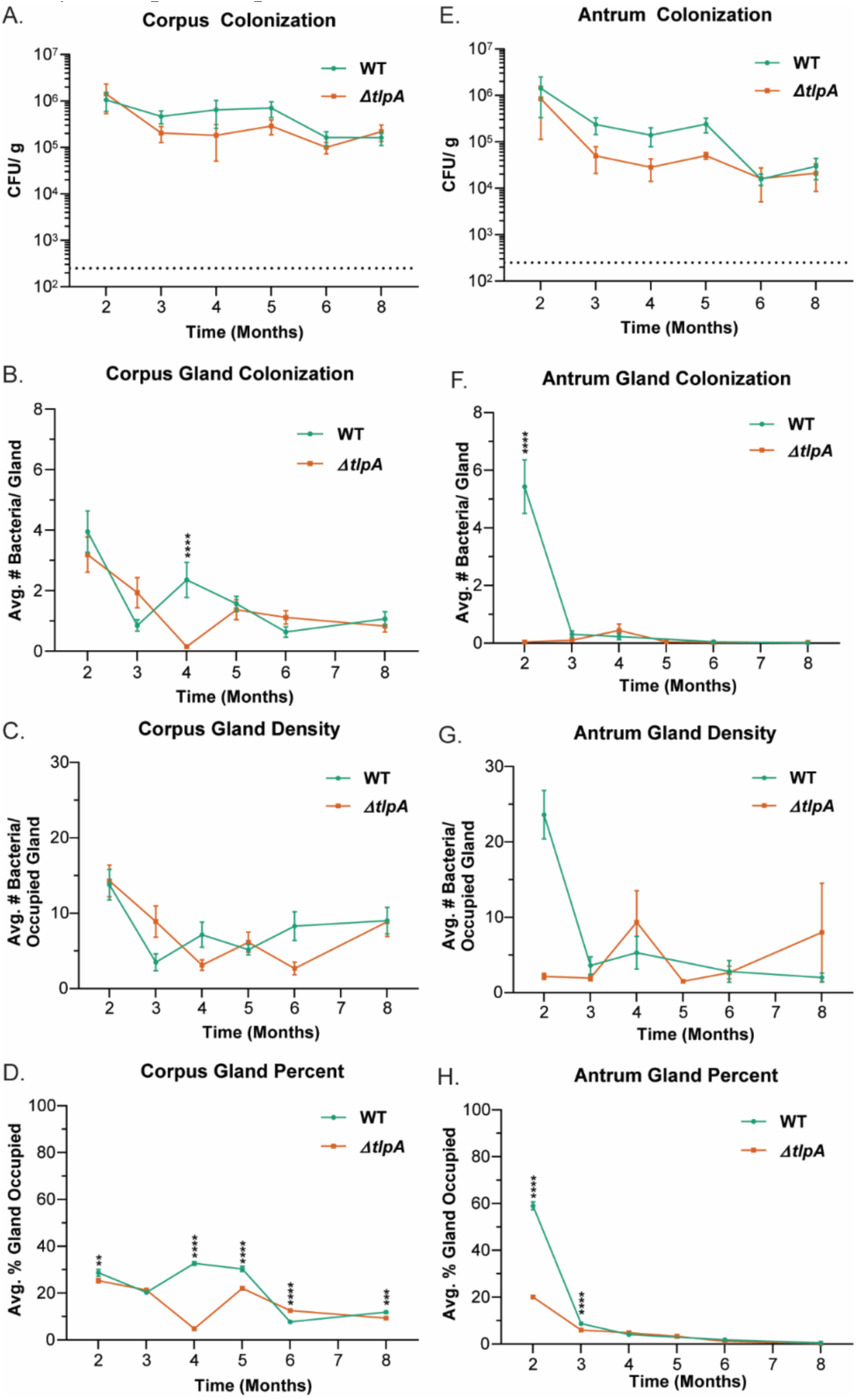
A replicate time course infection confirms few differences in gland colonization between WT and Δ*tlpA* SS1 during the chronic stage of infection. Female C57BL/6N mice (n ≥ 3) were orally infected with WT SS1 GFP^+^ or Δ*tlpA* SS1 GFP^+^ for 2-, 3-, 4-, 5-, 6-, and 8-months. Colonization of WT SS1 and Δ*tlpA* in the corpus was assessed for (A) total colonization, (B) gland colonization, (C) gland density, and (D) gland percent. Colonization of WT SS1 and Δ*tlpA* in the antrum was assessed for (E) total colonization, (F) gland colonization, (G) gland density, and (H) gland percent. For all graphs, error bars represent the standard error of the mean. *, P<0.05; **, P<0.01; ***, P<0.001; ****, P<0.0001, comparisons between strains using two-way ANOVA, Šídák multiple comparison test.

**Supplemental Figure 6.**
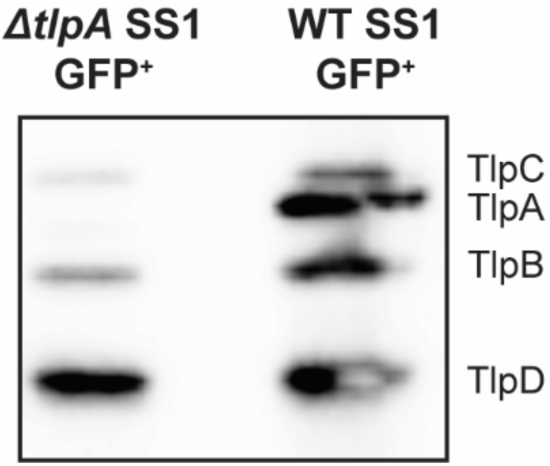
Verification of Δ*tlpA* SS1 GFP^+^. Verification of the generation of a Δ*tlpA* clean deletion mutant in a GFP^+^ SS1 background was confirmed via PCR (Data not shown) and western blot. Anti-TlpA22 (33) was used to detect the conserved, cytoplasmic portion of *H. pylori* TlpA, B, C, and D. All Δ*tlpA* SS1 GFP^+^ clones lack TlpA but express all other chemoreceptors.

## REFERENCES

1. Harris PR, Smythies LE, Smith PD, Perez-Perez GI. 2013. Role of childhood infection in the sequelae of *H. pylori* disease. Gut Microbes 4:426–438.

2. Salama NR, Hartung ML, Müller A. 2013. Life in the human stomach: persistence strategies of the bacterial pathogen Helicobacter pylori. Nat Rev Microbiol 11:385–399.

3. Cover TL, Blaser MJ. 2009. Helicobacter pylori in Health and Disease. Gastroenterology 136:1863–1873.

4. Arnold IC, Dehzad N, Reuter S, Martin H, Becher B, Taube C, Müller A. 2011. *Helicobacter pylori* infection prevents allergic asthma in mouse models through the induction of regulatory T cells. J Clin Invest 121:3088–3093.

5. Ferlay J, Soerjomataram I, Dikshit R, Eser S, Mathers C, Rebelo M, Parkin DM, Forman D, Bray F. 2015. Cancer incidence and mortality worldwide: Sources, methods and major patterns in GLOBOCAN 2012. Int J Cancer 136:E359–E386.

6. Hooi JKY, Lai WY, Ng WK, Suen MMY, Underwood FE, Tanyingoh D, Malfertheiner P, Graham DY, Wong VWS, Wu JCY, Chan FKL, Sung JJY, Kaplan GG, Ng SC. 2017. Global Prevalence of Helicobacter pylori Infection: Systematic Review and Meta-Analysis. Gastroenterology 153:420–429.

7. Plummer M, Franceschi S, Vignat J, Forman D, de Martel C. 2015. Global burden of gastric cancer attributable to Helicobacter pylori. Int J Cancer J Int Cancer 136:487–490.

8. Howitt MR, Lee JY, Lertsethtakarn P, Vogelmann R, Joubert L-M, Ottemann KM, Amieva MR. 2011. ChePep Controls Helicobacter pylori Infection of the Gastric Glands and Chemotaxis in the Epsilonproteobacteria. mBio 2:e00098–11.

9. Keilberg D, Ottemann KM. 2016. How *H elicobacter pylori* senses, targets and interacts with the gastric epithelium: *H. pylori* -gastric epithelium interaction. Environ Microbiol 18:791–806.

10. Schreiber S, Konradt M, Groll C, Scheid P, Hanauer G, Werling H-O, Josenhans C, Suerbaum S. 2004. The spatial orientation of Helicobacter pylori in the gastric mucus. Proc Natl Acad Sci 101:5024–5029.

11. Yang C, Ottemann KM. 2019. Control of bacterial colonization in the glands and crypts. Curr Opin Microbiol 47:38–44.

12. Schreiber S, Bucker R, Groll C, Azevedo-Vethacke M, Garten D, Scheid P, Friedrich S, Gatermann S, Josenhans C, Suerbaum S. 2005. Rapid Loss of Motility of Helicobacter pylori in the Gastric Lumen In Vivo. Infect Immun 73:1584–1589.

13. Goyal RK, Guo Y, Mashimo H. 2019. Advances in the physiology of gastric emptying. Neurogastroenterol Motil 31:e13546.

14. Atuma C, Strugala V, Allen A, Holm L. 2001. The adherent gastrointestinal mucus gel layer: thickness and physical state in vivo. Am J Physiol Gastrointest Liver Physiol 280:G922–G929.

15. Schreiber S, Scheid P. 1997. Gastric mucus of the guinea pig: proton carrier and diffusion barrier. Am J Physiol-Gastrointest Liver Physiol 272:G63–G70.

16. Creamer B, Shorter RG, Bamforth J. 1961. The turnover and shedding of epithelial cells: Part I The turnover in the gastro-intestinal tract. Gut 2:110–116.

17. Sigal M, Rothenberg ME, Logan CY, Lee JY, Honaker RW, Cooper RL, Passarelli B, Camorlinga M, Bouley DM, Alvarez G, Nusse R, Torres J, Amieva MR. 2015. Helicobacter pylori Activates and Expands Lgr5+ Stem Cells Through Direct Colonization of the Gastric Glands. Gastroenterology 148:1392–1404.e21.

18. Zhang X. 2020. Mechanisms of persistence, innate immune activation and immunomodulation by the gastric pathogen Helicobacter pylori. Curr Opin Microbiol 10.

19. Johnson KS, Ottemann KM. 2018. Colonization, localization, and inflammation: the roles of H. pylori chemotaxis in vivo. Curr Opin Microbiol 41:51–57.

20. Lertsethtakarn P, Ottemann KM, Hendrixson DR. 2011. Motility and Chemotaxis in *Campylobacter* and *Helicobacter*. Annu Rev Microbiol 65:389–410.

21. Cerda O, Rivas A, Toledo H. 2003. Helicobacter pylori strain ATCC700392 encodes a methyl-accepting chemotaxis receptor protein (MCP) for arginine and sodium bicarbonate. FEMS Microbiol Lett 224:175–181.

22. Cerda OA, Núñez-Villena F, Soto SE, Ugalde JM, López-Solís R, Toledo H. 2011. tlpA gene expression is required for arginine and bicarbonate chemotaxis in Helicobacter pylori. Biol Res 44:277–282.

23. Huang JY, Sweeney EG, Sigal M, Zhang HC, Remington SJ, Cantrell MA, Kuo CJ, Guillemin K, Amieva MR. 2015. Chemodetection and Destruction of Host Urea Allows Helicobacter pylori to Locate the Epithelium. Cell Host Microbe 18:147–156.

24. Machuca MA, Johnson KS, Liu YC, Steer DL, Ottemann KM, Roujeinikova A. 2017. Helicobacter pylori chemoreceptor TlpC mediates chemotaxis to lactate. Sci Rep 7:14089.

25. Johnson KS, Elgamoudi BA, Jen FE-C, Day CJ, Sweeney EG, Pryce ML, Guillemin K, Haselhorst T, Korolik V, Ottemann KM. 2021. The dCache Chemoreceptor TlpA of *Helicobacter pylori* Binds Multiple Attractant and Antagonistic Ligands via Distinct Sites. mBio 12.

26. Collins KD, Andermann TM, Draper J, Sanders L, Williams SM, Araghi C, Ottemann KM. 2016. The Helicobacter pylori CZB Cytoplasmic Chemoreceptor TlpD Forms an Autonomous Polar Chemotaxis Signaling Complex That Mediates a Tactic Response to Oxidative Stress. J Bacteriol 198:1563–1575.

27. Croxen MA, Sisson G, Melano R, Hoffman PS. 2006. The Helicobacter pylori Chemotaxis Receptor TlpB (HP0103) Is Required for pH Taxis and for Colonization of the Gastric Mucosa. J Bacteriol 188:2656–2665.

28. Huang JY, Goers Sweeney E, Guillemin K, Amieva MR. 2017. Multiple Acid Sensors Control Helicobacter pylori Colonization of the Stomach. PLOS Pathog 13:e1006118.

29. Perkins A, Tudorica DA, Amieva MR, Remington SJ, Guillemin K. 2019. Helicobacter pylori senses bleach (HOCl) as a chemoattractant using a cytosolic chemoreceptor. PLOS Biol 17:e3000395.

30. Schweinitzer T, Mizote T, Ishikawa N, Dudnik A, Inatsu S, Schreiber S, Suerbaum S, Aizawa S-I, Josenhans C. 2008. Functional Characterization and Mutagenesis of the Proposed Behavioral Sensor TlpD of Helicobacter pylori. J Bacteriol 190:3244–3255.

31. Collins KD, Hu S, Grasberger H, Kao JY, Ottemann KM. 2018. Chemotaxis Allows Bacteria To Overcome Host-Generated Reactive Oxygen Species That Constrain Gland Colonization. Infect Immun 86:e00878–17.

32. Keilberg D, Zavros Y, Shepherd B, Salama NR, Ottemann KM. 2016. Spatial and Temporal Shifts in Bacterial Biogeography and Gland Occupation during the Development of a Chronic Infection. mBio 7:e01705–16, /mbio/7/5/e01705-16.atom.

33. Williams SM, Chen Y-T, Andermann TM, Carter JE, McGee DJ, Ottemann KM. 2007. Helicobacter pylori chemotaxis modulates inflammation and bacterium-gastric epithelium interactions in infected mice. Infect Immun 75:3747–3757.

34. Andermann TM, Chen Y-T, Ottemann KM. 2002. Two Predicted Chemoreceptors of *Helicobacter pylori* Promote Stomach Infection. Infect Immun 70:5877–5881.

35. Rolig AS, Shanks J, Carter JE, Ottemann KM. 2012. Helicobacter pylori Requires TlpD-Driven Chemotaxis To Proliferate in the Antrum. Infect Immun 80:3713–3720.

36. Arnold IC, Lee JY, Amieva MR, Roers A, Flavell RA, Sparwasser T, Müller A. 2011. Tolerance Rather Than Immunity Protects From *Helicobacter pylori* –Induced Gastric Preneoplasia. Gastroenterology 140:199–209.e8.

37. White JR, Winter JA, Robinson K. 2015. Differential inflammatory response to Helicobacter pylori infection: Etiology and clinical outcomes. J Inflamm Res 8:137–147.

38. Wroblewski LE, Peek RM, Wilson KT. 2010. Helicobacter pylori and Gastric Cancer: Factors That Modulate Disease Risk. Clin Microbiol Rev 23:713–739.

39. Arnold IC, Artola-Borán M, Tallón de Lara P, Kyburz A, Taube C, Ottemann K, van den Broek M, Yousefi S, Simon H-U, Müller A. 2018. Eosinophils suppress Th1 responses and restrict bacterially induced gastrointestinal inflammation. J Exp Med 215:2055–2072.

40. Arnold IC, Zhang X, Urban S, Artola-Borán M, Manz MG, Ottemann KM, Müller A. 2017. NLRP3 Controls the Development of Gastrointestinal CD11b + Dendritic Cells in the Steady State and during Chronic Bacterial Infection. Cell Rep 21:3860–3872.

41. Oertli M, Sundquist M, Hitzler I, Engler DB, Arnold IC, Reuter S, Maxeiner J, Hansson M, Taube C, Quiding-Järbrink M, Müller A. 2012. DC-derived IL-18 drives Treg differentiation, murine Helicobacter pylori–specific immune tolerance, and asthma protection. J Clin Invest 122:1082–1096.

42. Lundgren A, Trollmo C, Edebo A, Svennerholm A-M, Lundin BS. 2005. Helicobacter pylori-Specific CD4ϩ T Cells Home to and Accumulate in the Human Helicobacter pylori-Infected Gastric Mucosa. INFECT IMMUN 73:8.

43. Bamford KB, Fan X, Crowe SE, Leary JF, Gourley WK, Luthra GK, Brooks EG, Graham DY, Reyes VE, Ernst PB. 1998. Lymphocytes in the human gastric mucosa during Helicobacter pylori have a T helper cell 1 phenotype. Gastroenterology 114:482–492.

44. Eaton KA, Mefford M, Thevenot T. 2001. The Role of T Cell Subsets and Cytokines in the Pathogenesis of *Helicobacter pylori* Gastritis in Mice. J Immunol 166:7456–7461.

45. Gray BM, Fontaine CA, Poe SA, Eaton KA. 2013. Complex T Cell Interactions Contribute to Helicobacter pylori Gastritis in Mice. Infect Immun 81:740–752.

46. Rolig AS, Carter JE, Ottemann KM. 2011. Bacterial chemotaxis modulates host cell apoptosis to establish a T-helper cell, type 17 (Th17)-dominant immune response in Helicobacter pylori infection. Proc Natl Acad Sci 108:19749–19754.

47. Fung C, Tan S, Nakajima M, Skoog EC, Camarillo-Guerrero LF, Klein JA, Lawley TD, Solnick JV, Fukami T, Amieva MR. 2019. High-resolution mapping reveals that microniches in the gastric glands control Helicobacter pylori colonization of the stomach. PLOS Biol 17:e3000231.

48. Terry K, Williams SM, Connolly L, Ottemann KM. 2005. Chemotaxis Plays Multiple Roles during Helicobacter pylori Animal Infection. Infect Immun 73:803–811.

49. Carbo A, Olivares-Villagómez D, Hontecillas R, Bassaganya-Riera J, Chaturvedi R, Piazuelo MB, Delgado A, Washington MK, Wilson KT, Algood HMS. 2014. Systems Modeling of the Role of Interleukin-21 in the Maintenance of Effector CD4 ^+^ T Cell Responses during Chronic Helicobacter pylori Infection. mBio 5.

50. Martinez LE, O’Brien VP, Leverich C, Knoblaugh SE, Salama NR. 2018. Non-helical Helicobacter pylori show altered gland colonization and elicit less gastric pathology during chronic infection. preprint. Microbiology.

51. Dixon BREA, Hossain R, Patel RV, Algood HMS. 2019. Th17 Cells in *Helicobacter pylori* Infection: a Dichotomy of Help and Harm. Infect Immun 87:e00363–19, /iai/87/11/IAI.00363-19.atom.

52. Serrano C, Wright SW, Bimczok D, Shaffer CL, Cover TL, Venegas A, Salazar MG, Smythies LE, Harris PR, Smith PD. 2013. Downregulated Th17 responses are associated with reduced gastritis in Helicobacter pylori–infected children. Mucosal Immunol 6:950–959.

53. Shiomi S, Toriie A, Imamura S, Konishi H, Mitsufuji S, Iwakura Y, Yamaoka Y, Ota H, Yamamoto T, Imanishi J, Kita M. 2008. IL-17 is Involved in *Helicobacter pylori* -Induced Gastric Inflammatory Responses in a Mouse Model. Helicobacter 13:518–524.

54. Wagner A, Wang C, Fessler J, DeTomaso D, Avila-Pacheco J, Kaminski J, Zaghouani S, Christian E, Thakore P, Schellhaass B, Akama-Garren E, Pierce K, Singh V, Ron-Harel N, Douglas VP, Bod L, Schnell A, Puleston D, Sobel RA, Haigis M, Pearce EL, Soleimani M, Clish C, Regev A, Kuchroo VK, Yosef N. 2021. Metabolic modeling of single Th17 cells reveals regulators of autoimmunity. Cell S0092867421007005.

55. Thompson LJ, Danon SJ, Wilson JE, O’Rourke JL, Salama NR, Falkow S, Mitchell H, Lee A. 2004. Chronic *Helicobacter pylori* Infection with Sydney Strain 1 and a Newly Identified Mouse-Adapted Strain (Sydney Strain 2000) in C57BL/6 and BALB/c Mice. Infect Immun 72:4668–4679.

56. Algood HMS, Cover TL. 2006. *Helicobacter pylori* Persistence: an Overview of Interactions between *H. pylori* and Host Immune Defenses. Clin Microbiol Rev 19:597–613.

57. Chen W, Shu D, Chadwick VS. 2001. Reduced colonization of gastric mucosa by Helicobacter pylori in mice deficient in interleukin-101. J Gastroenterol Hepatol 16:377–383.

58. Ailloud F, Didelot X, Woltemate S, Pfaffinger G, Overmann J, Bader RC, Schulz C, Malfertheiner P, Suerbaum S. 2019. Within-host evolution of Helicobacter pylori shaped by niche-specific adaptation, intragastric migrations and selective sweeps. Nat Commun 10:2273.

59. Lee A, O’Rourke J, De Ungria MC, Robertson B, Daskalopoulos G, Dixon MF. 1997. A standardized mouse model of Helicobacter pylori infection: introducing the Sydney strain. Gastroenterology 112:1386–1397.

60. Rader BA, Wreden C, Hicks KG, Sweeney EG, Ottemann KM, Guillemin K. 2011. Helicobacter pylori perceives the quorum-sensing molecule AI-2 as a chemorepellent via the chemoreceptor TlpB. Microbiology 157:2445–2455.

61. Eaton KA, Radin MJ, Krakowka S. 1995. An Animal Model of Gastric Ulcer Due to Bacterial Gastritis in Mice. Vet Pathol 32:489–497.

62. Rugge M, Correa P, Dixon MF, Fiocca R, Hattori T, Lechago J, Leandro G, Price AB, Sipponen P, Solcia E, Watanabe H, Genta RM. 2002. Gastric mucosal atrophy: interobserver consistency using new criteria for classification and grading. Aliment Pharmacol Ther 16:1249–1259.

63. Arnold IC, Zhang X, Artola-Boran M, Fallegger A, Sander P, Johansen P, Müller A. 2019. BATF3-dependent dendritic cells drive both effector and regulatory T-cell responses in bacterially infected tissues. PLOS Pathog 15:e1007866.

